# Notch signaling restricts FGF pathway activation in parapineal cells to promote their collective migration

**DOI:** 10.1101/570820

**Authors:** Lu Wei, Patrick Blader, Myriam Roussigné

## Abstract

Coordinated migration of cell collectives is important during embryonic development and relies on cells integrating multiple mechanical and chemical cues. Recently, we described that focal activation of the FGF pathway promotes the migration of the parapineal in the zebrafish epithalamus. How FGF activity is restricted to leading cells in this system is, however, unclear. Here, we address the role of Notch signaling in modulating FGF activity within the parapineal. While Notch loss-of-function results in an increased number of parapineal cells activating the FGF pathway, global activation of Notch signaling decreases it; both contexts result in defects in parapineal migration and specification. Decreasing or increasing FGF signaling in a Notch loss-of-function context respectively rescues or aggravates parapineal migration defects without affecting parapineal cells specification. We propose that Notch signaling controls the migration of the parapineal through its capacity to restrict FGF pathway activation to a few leading cells.

## Introduction

Coordinated migration of cell collectives is a widespread phenomenon, being seen predominantly during embryonic development but also during tissue repair in adults, for example. The molecular and cellular mechanisms underlying collective cell migration have been studied *in vivo* in different model organisms (Friedl and Gilmour, 2009; Ochoa-Espinosa and Affolter, 2012; Pocha and Montell, 2014; Theveneau and Mayor, 2013). Recent progress in the analysis of mechanical forces together with the development of *in vitro* models and *in silico* modelling have improved our understanding of coordinated cell migration. Such studies have highlighted the variability in mechanisms from one model to another, indicating that collective migration is a highly adaptive and plastic process (Haeger et al., 2015; Theveneau and Linker, 2017).

Members of the FGF family of secreted signals have been implicated in many models of cell migration. For example, FGF signaling is described to promote migration of cell collectives, potentially through chemotaxis (Kadam et al., 2012), through the modulation of cell adhesiveness (Ciruna et al., 1997; McMahon et al., 2010) or by increasing random cell motility (Benazeraf et al., 2010). In the lateral line primordium, the FGF pathway is required for Notch-dependent formation of neuromast rosettes at the trailing edge of the migrating primordium (Durdu et al., 2014; Kozlovskaja-Gumbrienė et al., 2017; Lecaudey et al., 2008; Nechiporuk and Raible, 2008) and to maintain cluster cohesion (Dalle Nogare et al., 2014), with both of these processes being required for proper lateral line primordium migration. Despite the widespread and iterative role of the FGF pathway in cell migration models, however, it’s not clear how the dynamics of FGF signaling correlate with cell behaviours and how this can be modulated by other signals.

The parapineal is a small group of cells that segregates from the anterior part of the pineal gland at the midline of zebrafish epithalamus and migrates in an FGF-dependent manner to the left side of the brain (Concha et al., 2000; Duboc et al., 2015; Roussigne et al., 2012). To characterize the dynamics of FGF pathway activation during parapineal migration, we recently analyzed the temporal and spatial activation of a previously described FGF pathway reporter transgene, *Tg(dusp6:d2GFP)* (Molina et al., 2007; Roussigné et al., 2018). Using this reporter, we showed that the FGF pathway is activated in an Fgf8-dependant manner in only a few parapineal cells located at the migration front and that experimentally activating the FGF pathway in a few parapineal cells restores parapineal migration in *fgf8-/-*mutant embryos. Taken together, these findings indicate that the restricted activation of FGF signaling in the parapineal promotes the migration of the parapineal cell collective. While the parapineal can receive Fgf8 signals from both sides of the midline, focal pathway activation is primarily detected on the left. This asymmetry in FGF pathway activation requires the TGFβ/Nodal signaling pathway, which is activated on the left side of the epithalamus prior to parapineal migration (Bisgrove et al., 1999; Concha et al., 2000; Liang et al., 2000). Although the Nodal pathway appears to bias the focal activation of FGF signaling to the left, after a significant delay the restriction of FGF activity still occurs in the absence of Nodal activity and the parapineal migrates (Roussigné et al., 2018).

All parapineal cells appear competent to activate the FGF pathway begging the question as to how the activation of the pathway is restricted to only a few cells. In this study, we address whether Notch signaling might modulate the activation of FGF pathway in the parapineal. We show that while loss-of-function of Notch leads to expanded FGF pathway activation in the parapineal, activating the Notch pathway causes a strong reduction in the expression of the FGF reporter transgene, with both contexts leading to defects in parapineal migration. Loss or gain of function for Notch signaling also interferes with the specification of parapineal cell identity; loss-of-function results in a significant increase in the number of *gfi1ab* and *sox1a* expressing parapineal cells, whereas gain of Notch activity results in the opposite phenotype. In contrast, the number of parapineal cells expressing *tbx2b*, a putative marker for parapineal progenitors cells (Snelson et al., 2008), is not affected in either loss or gain of function for Notch. Pharmacological inhibition of Notch pathway activity suggests that the roles of Notch in the specification and migration of parapineal cells can be uncoupled. Finally, a global decrease or an increase in the level of FGF signaling can respectively rescue or aggravate the parapineal migration defect caused by Notch loss-of-function but without affecting the specification of parapineal cells. Our data indicate that the Notch pathway regulates the specification and migration of parapineal cells independently and that the role of Notch signaling in promoting parapineal migration, but not specification, depends on its ability to restrict FGF pathway activation to a few parapineal cells.

## Results

### The parapineal of *mindbomb* mutant embryos display expanded FGF pathway activation

In models of cell migration during sprouting of tubular epithelia, Notch-Delta mediated cell-cell communication contributes to tip cell selection by restricting the ability of followers cells to activate RTK signaling (Ghabrial and Krasnow, 2006; Ikeya and Hayashi, 1999; Riahi et al., 2015; Siekmann and Lawson, 2007a). To address whether Notch signaling could similarly restrict FGF pathway activation in the freely moving group of parapineal cells, and thus promote its migration, we analyzed the expression of an FGF pathway activity reporter transgene, *Tg(dusp6:d2EGFP)* (Molina et al., 2007), in embryos mutant for the *mindbomb* (*mib*^*ta52b*^) gene, a well described loss-of-function context for the Notch pathway (Itoh et al., 2003). At 32 hours post-fertilisation (hpf), we observed a larger number of *Tg(dusp6:d2EGFP)* expressing cells in the parapineal of *mib*^−/−^ mutant embryos (7±3 d2EGFP+ cells) compared to siblings (4±2 d2EGFP+ cells; p-value=3.845e-05) (**Figures 1A-1B’, 1E**). The increase in *Tg(dusp6:d2EGFP)* expressing cells was not accompanied by an increase in the number of parapineal cells expressing *sox1a*, the earliest described parapineal specific marker (Clanton et al., 2013) (**Figures 1C-1D, 1G**), although we observed a slight increase in the total number of parapineal cells as determined using nuclear staining to visualize the parapineal rosette (22±7 in *mib*^−/−^ mutants compared to 19±5 in sibling control embryos; p-value=0.037) (**Figures 1F**).

**Figure 1.**
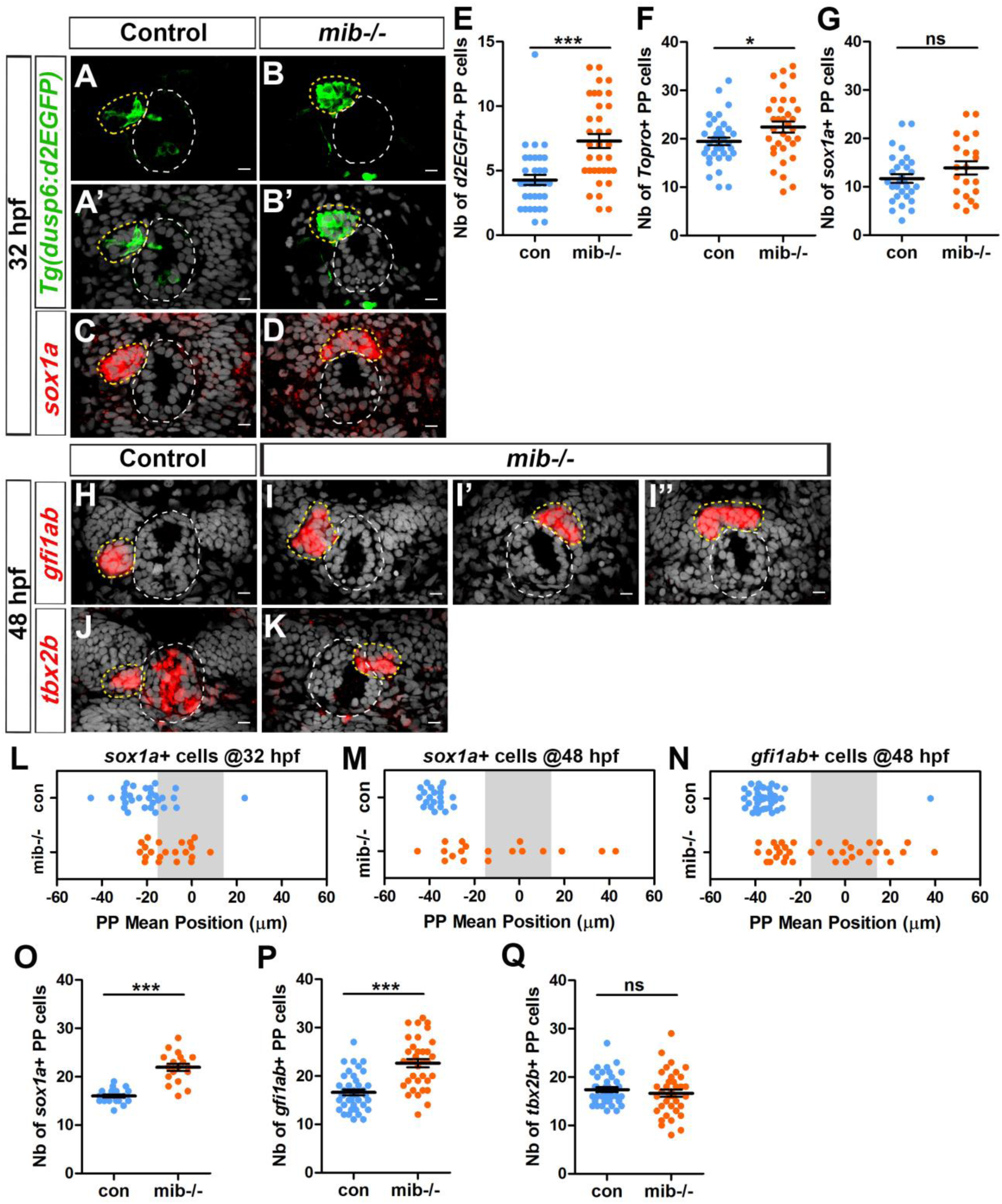
Increased activation of FGF signaling in *midbomb* mutants correlates with defects in migration and specification of parapineal cells. (A-D) Confocal maximum projection (A-B) or sections (A’-D) showing the expression of the *Tg(dusp6:d2EGFP)* transgene (Green, A-B’) or *sox1a* (red, C-D) in the epithalamia of 32 hpf in control (A, A’, n=36 and C, n=30) and in *mib*^−/−^ mutant embryos (B, B’, n=34 and D, n=21). (E-G) Dot plots showing the number (E-G) of *Tg(dusp6:d2EGFP)* (E), Topro-3 (F) and *sox1a* (G) positive parapineal cells in control (blue dots) or *mib*^−/−^ mutant embryos (orange dots) at 32 hpf, with mean ± SEM. *Tg(dusp6:d2EGFP)* FGF reporter is expressed in more parapineal cells in *mib*^−/−^ than controls (E; 7±3 d2EGFP+ cells in *mib*^−/−^ mutant compared 4±2 in siblings; p-value=3.845e-05, Welch t-test) while the expression of *sox1a* is similar in both contexts (G); the average number of parapineal cells counted with a nuclear marker increases slightly (F; 22±7 in *mib*^−/−^ mutants versus 19±5 in sibling control embryos; p-value=0.037, Welch t-test). (H-I’’) Confocal sections showing the expression of *gfi1ab* (red) at 48 hpf in control embryos (H; n=40) and in three examples of *mib*^−/−^ mutant embryos (I-I’’; n=35). (J-K) Confocal sections showing the expression of *tbx2b* (red) at 48 hpf in control embryos (J; n=39) and in one example of *mib*^−/−^ mutants embryo with the parapineal on the right (K; n=36). (L-N) Dot plots showing the mean position of parapineal cells expressing *sox1a* at 32 hpf (L) *sox1a* at 48 hpf (M) *or gfi1ab* (N) (in µm distance to the brain midline (x=0)) with each dot representing an embryo. Parapineal migration is usually delayed in *mib*^−/−^ mutants at 32 hpf (L). At 48 hpf, the parapineal of *mib*^−/−^ mutant embryos either did not migrate (n=12/35 with a parapineal mean position between −15µM and +15µM (shaded zone) relative to brain midline (Reference 0)) or migrates either to the left (n=17/35) or to the right (n=6/35) (N, orange dots), while it usually migrated to the left in control embryos (N, n=39/40, blue dots); p-value<0.0001, Welch t-test on absolute values. (O-Q) Number of parapineal cells expressing *sox1a* (O), *gfi1ab* (P) and *tbx2b* (Q) at 48 hpf in control (blue dots) or in *mib*^−/−^ mutant embryos (orange dots). The number of *sox1a* and *gfi1ab+* positive parapineal cells at 48 hpf is increased in *mib*^−/−^ mutant embryos (p-value<0.0001 in Welch t-test) compared with controls (O, P) while the number of *tbx2b* expressing cells is unchanged (Q). Confocal sections are merged with a nuclear staining (grey) that makes the epiphysis (white circle) and parapineal (yellow circle) visible. Embryo view is dorsal, anterior is up; scale bar=10 µm. Mean ± SEM are indicated as long and short bars. *** p-value<0.0001; * p-value<0.05 in Welch t-test. Data are representative of three experiments (H-I’’, N, P) or two independent experiments (A-G, J-M, O, Q). See also Figure 1–figure supplement 1.

To confirm that the phenotypes observed in *mib*^−/−^ mutants are caused by Notch pathway loss-of-function, we analyzed the parapineal in embryos injected with a morpholino (MO; antisense oligonucleotide) blocking translation of the two zebrafish orthologs of *su(H)/rbpj* (Echeverri and Oates, 2007); Rpbj proteins are transcription factors required for canonical Notch signaling (Fortini and Artavanis-Tsakonas, 1994; Hsieh et al., 1996). As observed in *mib*^−/−^ mutants, the proportion of *Tg(dusp6:d2EGFP)* expressing cells was increased in the parapineal of embryos injected with *rbpj a/b* MO (4ng) (10±6 d2EGFP+ cells) compared to controls (6±4 d2EGFP+ cells; p-value=0,0016) (**Figure 1–figure supplement 1, C**). Altogether, our results indicate that the FGF signaling pathway is activated in more parapineal cells when Notch signaling is abrogated.

### Loss of Notch signaling results in defects in parapineal migration

Interestingly, while in controls the parapineal had usually initiated migration at 32 hpf, it was still detected at the midline in most stage-matched *mib*^−/−^ mutant embryos (**Figures 1A-D, 1L**). To address the effect of Notch inhibition on parapineal migration further, we examined *mib*^−/−^ mutant embryos at a later stage, when the parapineal had unambiguously migrated in all controls. Analyzing *sox1a* (Clanton et al., 2013) or *gfi1ab* (Dufourcq et al., 2004) expression at 2 days post-fertilization revealed that the parapineal failed to migrate in about a third of *mib*^−/−^ embryos (defined by parapineal mean position within −15 µm and +15 µm of the midline (grey shaded zone)) (**Figures 1M, 1N**). Using *gfi1ab* as a marker, for example, we found that the parapineal had not migrated in 35% of *mib*^−/−^ embryos (n=12 of 35 embryos; **Figures 1I”, 1N**), while it migrated to the left in more than 95% of control embryos (n=39/40, **Figures 1H, 1N**, p-value<0.0001); the parapineal in *mib*^−/−^ mutant embryos also migrated to the right more often than in controls (17%, n=6/35; **Figures 1I’, 1N**).

In embryos injected with *rbpj a/b* MO (4ng), the mean position of parapineal cells at 32 hpf was closer to the midline, suggesting a similar defect in parapineal migration as observed in *mib*^−/−^ mutant embryos (**Figure 1–figure supplement 1, D).** Using *gfi1ab* as a marker at 48 hpf, we found that both parapineal migration *per se* and the orientation of migration were significantly affected in *rbpj a/b* morphants (**Figure 1–figure supplement 1, A-B’, E**). While the parapineal migrated to the left in most uninjected controls (94%, n=31/33), in *rbpj a/b* morphants migration was blocked (13% of the embryos, n=6/46; p-value=0.0001) or its orientation was partially randomized (left in 67% of the embryos, n=31/46; right in 20% of the embryos, n=9/46; p-value=0.0002) (**Figure 1–figure supplement 1, A-B’, E**).

Taken together, our data show that loss of Notch signaling results in defects both in parapineal migration *per se* (distance from the midline) and in the laterality of migration (left orientation), and that this correlates with the FGF signaling pathway being activated in more parapineal cells.

### Loss of Notch signaling results in an increase in the number of certain parapineal cell subtypes

While quantifying parapineal mean position at 48 hpf, we observed an overall increased in number of *sox1a* expressing cells in *mib*^−/−^ mutants at this stage (**Figure 1O**) despite finding no significant change at 32 hpf (**Figure 1G**). Similarly, we found that the number of *gfi1ab* positive cells at 48 hpf was significantly increased in *mib*^−/−^ mutants (23±5) compared to control embryos (17±4; p-value=3.901e-07) (**Figures 1H-1I’’, 1P**). In embryos injected with 4 ng of *rbpj a/b* MO, we also observed that the number of *gfi1ab* expressing parapineal cells at 48 hpf was significantly increased (19±6) compared to control embryos (14±3; p-value<0.0001) (**Figure 1– figure supplement 1, A-B’, F**). In contrast, the number of parapineal cells expressing *tbx2b*, a marker previously suggested to label parapineal progenitors (Snelson et al., 2008), was not increased in *mib*^−/−^ mutants (**Figure 1J-1K, 1Q**).

Taken together, our data show that loss of Notch signaling results in defects in parapineal migration and, at later stages, an increase in the number of *gfi1ab and sox1a* expressing parapineal cells (putative differentiated parapineal cells) while the number of *tbx2b* expressing cells (putative parapineal progenitors) is unchanged.

### The roles of Notch signaling in the specification and migration of parapineal cells can be uncoupled

Our data show that blocking Notch signaling leads to an expansion of FGF pathway activation in the parapineal, defects in parapineal migration and laterality, and an increase in the number of *gfi1ab* and *sox1a* expressing parapineal cells. With the aim of unraveling potential causative links between these different phenotypes, we used a pharmacological inhibitor of the γ-secretase complex, to block Notch signaling pathway during different time windows (Romero-Carvajal et al., 2015; Rothenaigner et al., 2011); γ-secretase activity is required for the release of the intracellular domain of Notch, NICD, during activation of the canonical pathway (Geling et al., 2002).

We first treated wild-type embryos with LY411575 between 22 and 32 hpf, a time window corresponding to parapineal segregation from the pineal gland and the onset of its migration. While no change in *gfi1ab* expression was detected in embryos treated with 30 µM LY411575 (**Figures 2C**), a higher concentration of LY411575 (100 µM) resulted in an increase in the number of *gfi1ab* positive cells in treated embryos (**Figures 2A-2C**); neither treatment resulted in defects in parapineal migration (**Figures 2D**). As this effect of LY411575 treatment was modest, we next treated embryos heterozygous for *mib* mutation (*mib*^+/−^) during the same time window thinking that this might provide a sensitized background for the drug. *mib*^+/−^ heterozygous embryos treated with the lower dose of LY411575 show a strong increase in the number of *gfi1ab* expressing parapineal cells (28±5) compared to LY411575 treated wild-type controls (17±3; p-value=1.0e-10) or DMSO treated *mib*^+/−^ heterozygotes (20±2; p-value=2.2e-10) (**Figures 2E-2H, 2I**). As before, however, we did not detect a parapineal migration defect in LY411575 treated *mib*^+/−^ embryos (**Figures 2E-2H, 2J**), even when we increased the dose of LY411575 to 200 µM (**Figure 2–figure supplement 1**). Therefore, although LY411575 treatment from 22 to 32hpf can synergize with a *mib*^+/−^ heterozygous genetic background to promote an increase in the number of *gfi1ab* positive parapineal cells, it does not affect parapineal migration. These data indicate that the role of Notch in controlling the specification of parapineal cells can be uncoupled from its function in parapineal migration.

**Figure 2.**
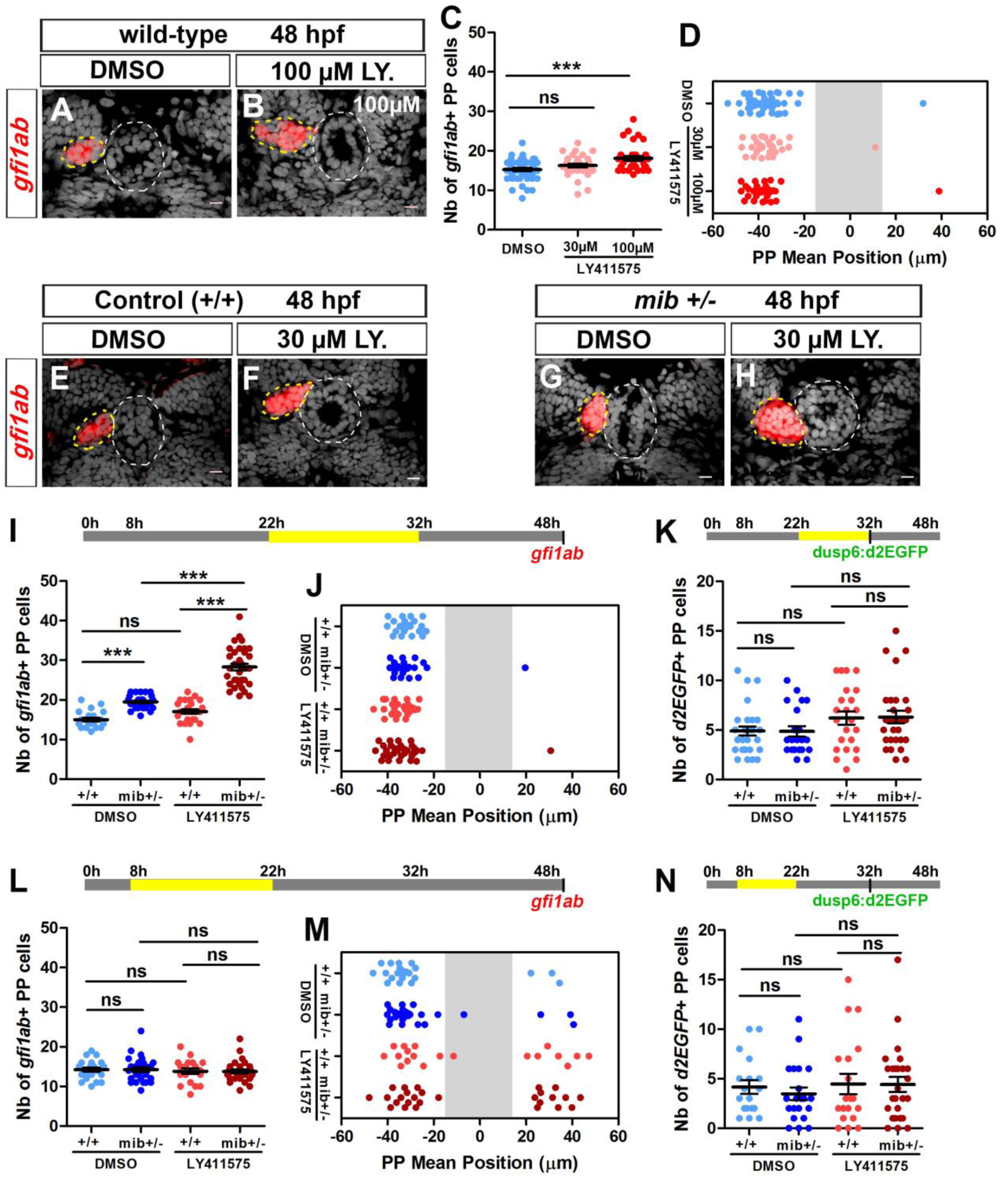
Uncoupled roles for the Notch pathway in the specification, laterality and migration of parapineal cells. (A-B) Confocal sections showing the expression of *gfi1ab* (red) in embryos treated with DMSO (A; n=24) or 100 µM LY411575 (B; n=32) from 22 to 32 hpf and fixed at 48 hpf, merged with nuclear staining (grey). Embryo view is dorsal, anterior is up; epiphysis (white circle) and parapineal (yellow circle); scale bar=10 µm. (C-D) Dot plots showing the number (C) and the mean position (D) of *gfi1ab* expressing parapineal (PP) cells at 48 hpf in embryos treated with DMSO (controls, blue dots, n=47), with 30 µM LY411575 (light red dots, n=31) or 100 µM LY411575 (red dots, n=32) from 22 hpf to 32 hpf, with mean ± SEM; *** p-value=0.0003. (E-H) Confocal sections showing the expression of *gfi1ab* (E-H) (red) merged with nuclear staining (grey) at 48 hpf, in wild-type (+/+) (E, F) or *mib*^+/−^ heterozygote embryos (F, H) treated with DMSO (E, n=23 or G, n=26) or with LY411575 (F, n=25 and G, n=34) from 22 to 32 hpf. (I-N) Upper panels show a schematic of the LY411575 treatment timeline, 22 to 32 hpf for I-K or 8 to 22 hfp for dot plots L-N. Dot plots showing the number (I, L) and the mean position (J, M) of *gfi1ab* expressing cells at 48 hpf, or the number of *Tg(dusp6:d2EGFP)* expressing cells at 32 hpf, in the parapineal of DMSO treated wild-type (+/+, dark blue dots; I-J, n=23; L-M, n=12; K, n=28; N, n=18), DMSO treated *mib*^+/−^ heterozygote (light blue dots; I-J, n=26; L-M, n=17; K, n=21; N, n=21), LY411575 treated wild-type (dark red dots; I-J, n=25; L-M, n=11; K, n=23, N, n=19) and LY411575 treated *mib*^+/−^ embryos (light red dots, I-J, n=34 or L-M, n=16; K, n=29, N, n=26); each dot represents a single embryo. Mean ± SEM is shown in I, K, L, N; *** p-value<0.0001, in Wilcoxon test and Welch t-test. J, M: there is no defect in migration *per se* (ns p-value in Welch t-test on absolute value) but LY411575 treatment from 8 to 22 hpf (M) triggers a laterality defect (increased number of embryos with a parapineal on the right). Data are representative of three (E-H, I-K) or two experiments (A-D and L-N). See also Figure 2–figure supplement 1 and Figure 2–figure supplement 2.

Finally, we saw no significant increase in the number of *Tg(dusp6:d2EGFP)* positive cells in *mib*^+/−^ heterozygous embryos treated with LY411575 during the 22 to 32 hpf time window (**Figures 2K**). This wild-type level of FGF pathway activation correlates with a correct migration of the parapineal in LY411575 treated embryos and is consistent with our previous results suggesting that restricted FGF pathway activation is important for correct parapineal migration (Roussigné et al., 2018).

### Notch activity is required early for unilateral activation of the Nodal pathway in the epithalamus

To determine whether the parapineal migration defects we observed might be an indirect consequence of an earlier role of Notch signaling, we also analyzed the epithalamus of embryos treated with LY411575 during an earlier 8 to 22 hpf time window. This early drug treatment did not interfere with the number of *gfi1ab* positive cells (**Figures 2L**), or with migration in itself (**Figure 2M**), but led to a partial randomization of the direction of parapineal migration as shown by a significant increase in the number of embryos with a right parapineal (**Figure 2M**).

We hypothesized that the partial randomization of parapineal sidedness observed in *mib*^−/−^ mutants, *rbpj a/b* morphants or in embryos treated with LY411575 from 8 to 22 hpf could be caused by changes in the activation pattern of Nodal signaling in the epithalamus (Concha et al., 2000; Liang et al., 2000; Regan et al., 2009). To address this possibility, we analyzed the expression of *pitx2c*, a Nodal signaling target gene (Concha et al., 2000; Essner et al., 2000; Liang et al., 2000), in the different contexts of Notch loss-of-function. While *pitx2c* expression is detected in the left epithalamus in control embryos between 28 and 32 hpf (n=26/28), we observed that its expression is bilateral in most *mib*^−/−^ mutant embryos (n=26/29) (**Figure 2– figure supplement 2, A-C, D),** in a majority of *rbpja/b* morphants (n=13/23, **Figure 2–figure supplement 2, E**) and in approximately half of the embryos treated with LY411575 from 8 to 22 hpf, regardless of whether they were heterozygotes for the *mib* mutation (*mib*^+/−^, n=6/10) or not (n=4/13) (**Figure 2–figure supplement 2, G**). In contrast, the expression of *pitx2c* was indistinguishable from controls in embryos treated with LY411575 during the later time window (22-32 hpf, **Figure 2–figure supplement 2, H**) or in embryos expressing NICD from 26 hpf (**Figure 2–figure supplement 2, F**), which was expected given the absence of laterality defects in these contexts.

Despite triggering defects in parapineal laterality, the early time window of LY411575 treatment (8 to 22 hpf) did not affect parapineal migration *per se* and, as observed for the late time window (22-32 hpf), this correlates with no significant change in the *Tg(dusp6:d2EGFP)* expression pattern (**Figure 2N**). Our data indicate that the partial randomization of parapineal migration observed in *mib*^−/−^ mutants or *rbpj* morphants is due to an early role of Notch pathway in restricting Nodal signaling to the left epithalamus and not to changes in the pattern of FGF activation.

### Notch gain of function inhibits FGF pathway activation in the parapineal, and blocks the specification and migration of parapineal cells

When we abrogated Notch signaling, we observed an increase in the number of *Tg(dusp6:d2EGFP)+* parapineal cells and correlated defects in parapineal migration (**Figures 1E, 1L-M, Figure 1–figure supplement 1, C-E**). To address further the role of the Notch pathway in modulating FGF activation, we analyzed the phenotypes associated with global activation of Notch signaling. For this, we used previously described transgenic lines, *Tg(hsp70:gal4)* and *Tg(UAS:NICD-myc)* (Scheer and Campos-Ortega, 1999), to induce widespread expression of the Notch Intracellular Domain (NICD) upon heat shock. In most embryos globally expressing NICD from 26 hpf, we observed a strong decrease in the number of *Tg(dusp6:d2EGFP)* expressing parapineal cells at 36 hpf (2.5±2 cells) compared with the control embryos (5±4; p-value=0.01) (**Figures 3A-3B, 3G**); the mean intensity of d2EGFP fluorescence was also significantly decreased in these embryos compared with the controls (p-value=0.0001) (**Figures 3A-3B, 3H**).

**Figure 3.**
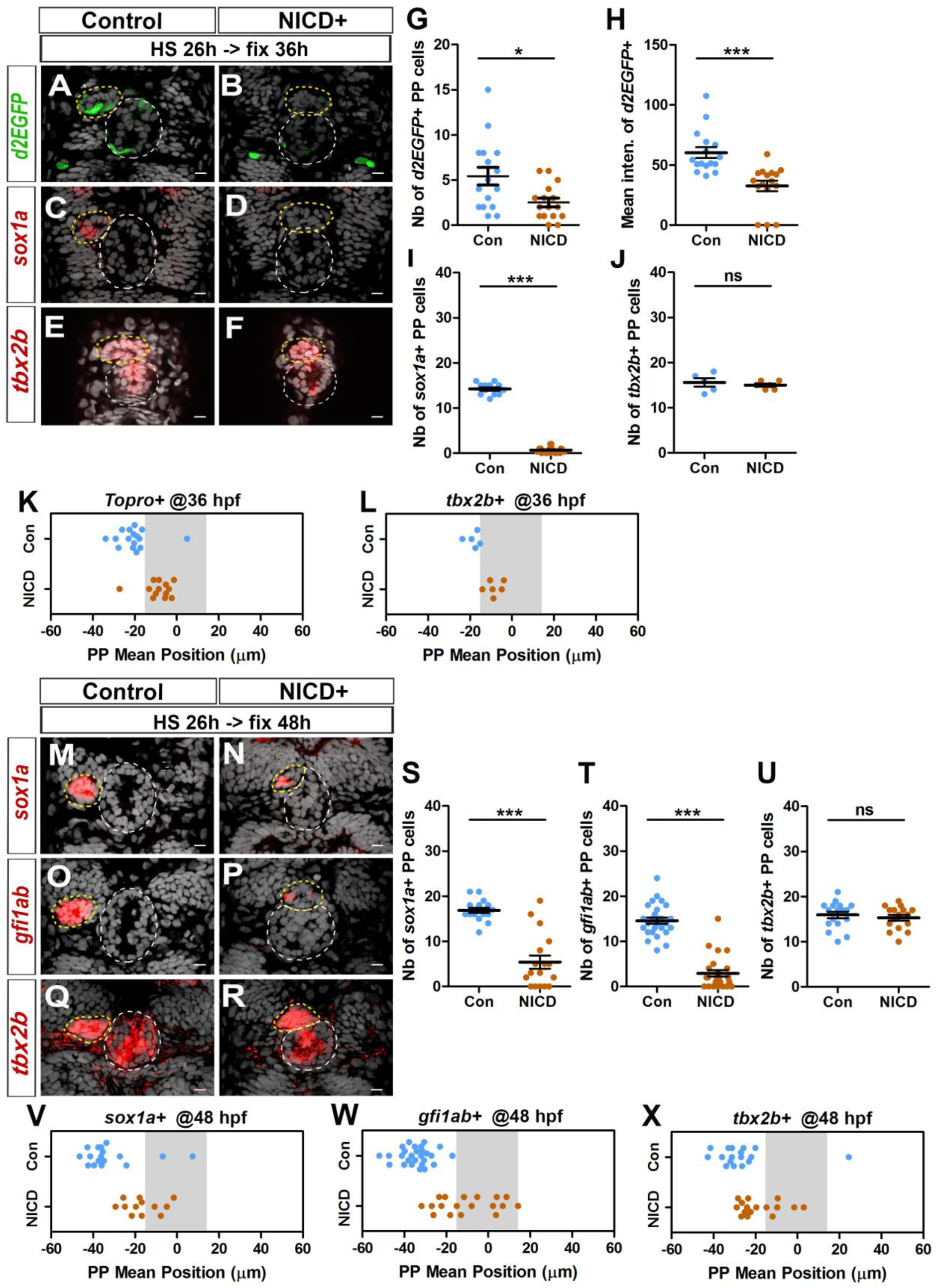
Ectopic Notch signaling triggers decreased FGF activation and defects in migration and specification of parapineal cells. (A-F) Confocal sections showing the expression of *Tg(dusp6:d2EGFP)* (A-B, green), *sox1a* (C-D, red) or *tbx2b* (E-F, red) at 36 hpf, in control embryos (A, n=16; C, n=12; E, n=5) or in *Tg(hsp70l:Gal4), Tg(UAS:myc-notch1a-intra)* embryos (B, n=16; D, n=17; F, n=6) following a heat-shock (HS) at 26 hpf; sections are merged with nuclei staining (grey). (G-J) Dot plots showing the number (G) and the mean intensity fluorescence (H) of *Tg(dusp6:d2EGFP)* expressing parapineal, or the number of *sox1a* (I) and *tbx2b* (J) expressing parapineal cells, in controls (blue dots) or in NICD expressing embryos (orange dots) at 36 hpf following heat shock at 26 hpf. (M-R) Confocal sections showing the expression of *sox1a* (M-N), *gfi1ab* (O-P) *or tbx2b* (Q-R) (red) merged with nuclei staining (grey), at 48 hpf, in the epithalamia of control (M, n=17; O, n=27; Q, n=17) or *Tg(hsp70l:Gal4);Tg(UAS:myc-notch1a-intra)* double transgenic embryos (N, n=17; P, n=25; R, n=16), following heat-shock (HS) at 26 hpf. The expression of *sox1a* and *gfi1ab* is lost or decreased while *tbx2b* expression is unchanged in the parapineal of NICD expressing embryos. (S-X) Dot plots showing the number of *sox1a* (S), *gfi1ab* (T) and *tbx2b* (U) expressing parapineal cells at 48 hpf in controls (blue dots) or in embryos expressing NICD after heat shock at 26 hpf (orange dots) and the corresponding mean position of the cells (W-X) when expression was detected (number of *sox1a*+ or *gfi1ab+* cells >0). In confocal sections, embryo view is dorsal, anterior is up; epiphysis (white circle) and parapineal gland (yellow circle); scale bar=10 µm. Mean ± SEM is indicated on dot plots G-J and S-U; *** p-value<0.0001, * p-value<0.05 in Welch t-test and Wilcoxon test.; For migration dot plots, p-value<0.01 (L) or p-value<0.001 (K and V-X) in pairwise Wilcoxon test and Welch t-test on absolute values. Data are representative of three (O, P, T, W) or two experiments (A-D, G-I, M-N, Q-R, S, U, V, X); data based on *tbx2b* expression at 36 hpf (E-F, J, L) represents one experiment.

To assess a potential correlation between the inhibition of FGF pathway activation and defects in parapineal migration, we sought to analyze the mean position of *sox1a* or *gfi1ab* expressing cells at 36 or 48 hpf as we had done for Notch loss-of-function embryos. However, following heat shock at 26 hpf, *sox1a* expression was lost in most 36 hpf embryos (**Figures 3C-D, 3I**) and was strongly decreased at 48 hpf (**Figures 3M-3N, 3S**). Similarly, although the number of *gfi1ab* positive cells did not vary significantly in *Tg(hsp70:gal4); Tg(UAS:NICD-myc)* embryos heat shocked at 22 and 24 hpf (**Figure 3–figure supplement 1, A-D, I-J**), it was strongly decreased in embryos expressing NICD beginning from 26, 28 and 32 hpf (**Figures 3O-P, 3T and Figure 3–figure supplement 1 E-H, K-L**); in embryos heat shocked at 26 hpf or 28 hpf, *gfi1ab* staining was often completely lost in the parapineal (n=7/25 or n=8/26 respectively) or detected in less than 4 cells (n=11/25 or n=14/26 respectively) (**Figures 3O-P, 3T and Figure 3–figure supplement 1, E-F, K**). However, nuclear staining indicated that the parapineal rosettes can be detected in most of the embryos expressing NICD (**Figure 3A-F, 3M-R**, yellow circle), suggesting that the parapineal does form despite global activation of Notch. Similarly, *tbx2b* expression was not affected in NICD expressing embryos either at 36 hpf (**Figure 3E-F, 3J**) or at 48 hpf (**Figure 3Q-R, 3U**). Thus, global Notch activation appears to inhibit the specification/differentiation of *gfi1ab+* or *sox1a*+ cells from *tbx2b* progenitors.

As NICD expression led to a loss of *Tg(dusp6:d2EGFP)* and *sox1a* expression, we first relied on nuclear staining to assess parapineal cells mean position in this context at 36 hpf. We found that the parapineal was closer to the midline in embryos expressing NICD from 26 hpf, (−8.7 ± 6.3 µM; n=16) compared to control embryos (−20.5 ± 8.3 µM; n=16; p-value=0.001) (**Figures 3A-F, 3K).** This migration defect was also revealed by analyzing the mean position of *tbx2b* expressing parapineal cells at 36 hpf (p-value=0.0017) (**Figure 3L).** Defects in parapineal migration were confirmed at 48 hpf using *sox1a, gfi1ab* and *tbx2b* as markers to assess PP mean position (**Figure 3M-R, 3V-X**); for instance, when *gfi1ab* positive cells were detected, their mean position was significantly closer to the midline in embryos expressing NICD from 26 hpf (−10.1 ± 14 µM; n=18) compared to control embryos (−34.6 ± 7.5 µM; n=27) (p-value<0.0001) (**Figures 3O-P, 3W)**. Parapineal mean position was also affected in embryos with NICD induced just before parapineal formation (heat shock at 22 and 24 hpf), or after parapineal formation (heat shock at 28 and 32 hpf), although in the latter case, the penetrance varies (**Figure 3– figure supplement 1, A-H, M-P**).

Altogether, our data show that global activation of Notch signaling inhibits the migration of parapineal cells, and that this is correlated with a decrease in the level of FGF signaling detected in the parapineal. Ectopic Notch signaling also decreases the number of differentiated parapineal cells, a phenotype opposite to that observed in Notch loss-of-function contexts.

### Decreasing FGF signaling rescues the parapineal migration defects in loss of Notch context while increasing FGF signaling aggravates it

Inhibiting or activating the Notch pathway results in reciprocal effects on FGF pathway activation in the parapineal, as seen by an increase or decrease in the number of *Tg(dusp6:d2EGFP)* expressing parapineal cells, respectively. Both contexts are also associated with defects in parapineal migration, suggesting that Notch-dependent control of FGF activation in parapineal cells is important for their collective migration. To investigate the link between Notch and FGF signaling further, we analyzed whether the migration phenotype observed in *mib*^−/−^ mutants could be rescued by decreasing FGF signaling, using a pharmacological inhibitor of the FGF pathway (Mohammadi et al., 1997). We have previously shown that treating wild-type embryos with 10 µM SU5402 interferes with parapineal migration (Regan et al., 2009) and with the expression of the *Tg(dusp6:d2EGFP)* FGF reporter (Roussigné et al., 2018). Treating embryos with 5 µM SU5402, however, does not affect parapineal migration (**Figures 4A-4B, 4F**); the parapineal migrates in all SU5402 treated embryos (n=31/31) as well as in all DMSO treated control embryos (n=32/32) (**Figure 4F,** purple versus blue dots). Using this suboptimal dose, parapineal migration was partially rescued in *mib*^−/−^ embryos, with 19% of SU5402 treated *mib*^−/−^ embryos showing mean parapineal position between −15 and +15 µm (n=8/41) compared to 52% of DMSO treated *mib*^−/−^ embryos (n=16/31) (**Figures 4C-4D, 4F,** yellow versus orange dots; p-value=0.01). Thus, decreasing the level of FGF signaling activity can partially restore parapineal migration in a context where FGF activation is expanded supporting the hypothesis that Notch promotes parapineal migration through restricting FGF pathway activation.

**Figure 4.**
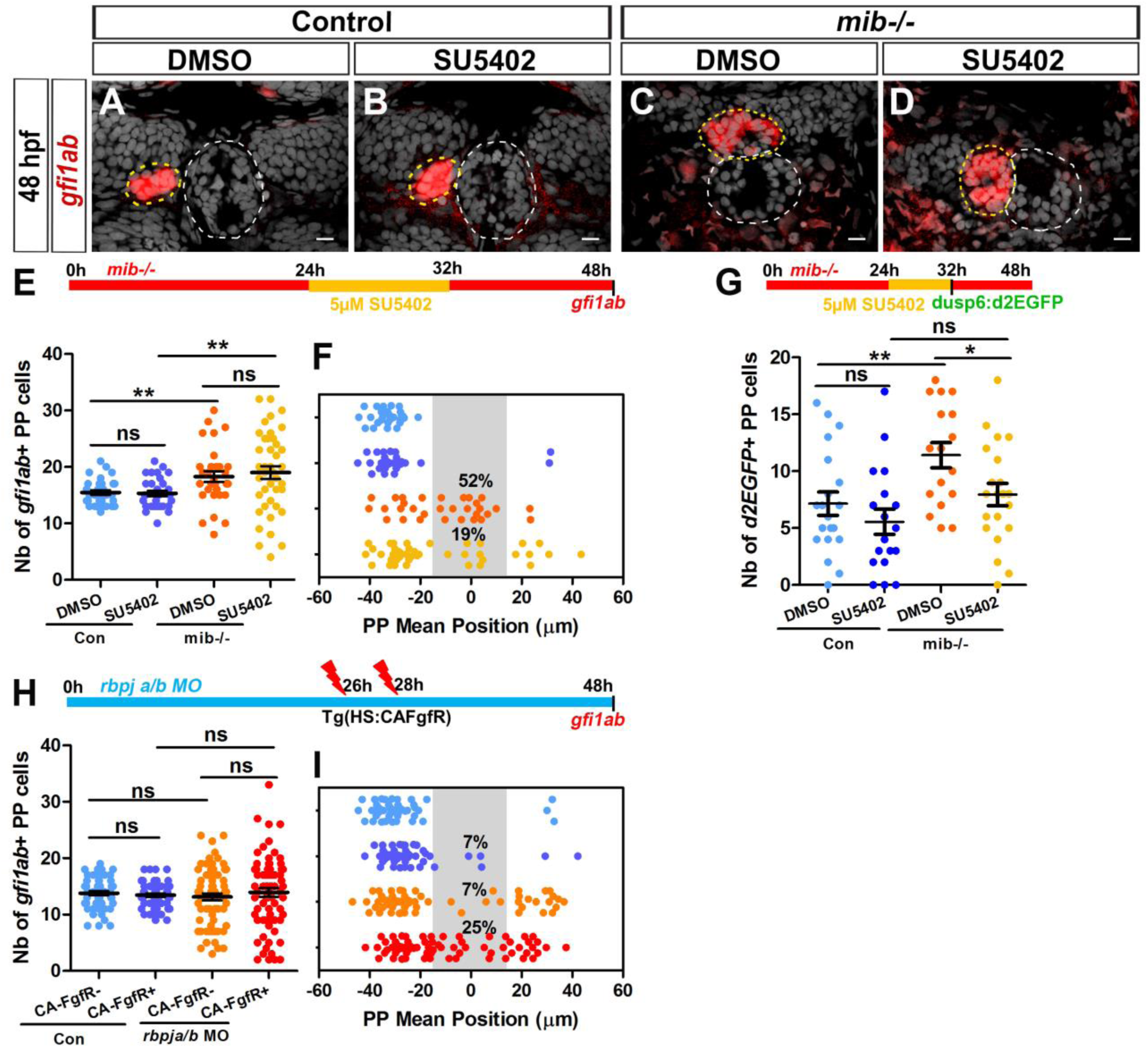
Decreasing or increasing FGF signaling rescues or aggravates the parapineal migration defect in Notch loss-of-function. (A-D) Confocal sections showing the expression of *gfi1ab* (red) merged with nuclei staining (grey) at 48 hpf in representative control sibling (A-B) or *mib*^−/−^ mutant embryos (C-D) treated from 24 to 32 hpf with DMSO (A, C) or 5 µM SU5402 (B, D). Parapineal migration is not affected in controls embryos treated with 5µM of SU5402 (A-B); in C-D, examples show a parapineal that failed to migrate in a DMSO treated *mib*^−/−^ mutant embryos (C) or that migrated to the left in a SU5402 treated *mib*^−/−^ mutant embryos (D). Embryo view is dorsal, anterior is up; epiphysis (white circle) and parapineal (yellow circle); scale bar=10 µm. (E-F) Upper panel show a schematic of the SU5402 (or DMSO) treatment timeline (24-32 hpf) in control and *mib*^−/−^ mutant embryos. Dot plots showing the number (E) and the mean position (F) of *gfi1ab* expressing parapineal (PP) cells at 48 hpf in control embryos treated with DMSO (light blue dots; n=32) or with 5µM SU5402 (dark blue dots, n=31), and in *mib*^−/−^ mutant embryos treated with DMSO (orange dots, n=31) or with SU5402 (yellow dots, n=41). The number of *gfi1ab* positive cells is increased in *mib*^−/−^ mutants embryos, regardless of whether they were treated with SU5402 or DMSO: DMSO control versus DMSO *mib*^−/−^, ** p-value=0.0042; SU5402 control versus SU5402 *mib*^−/−^, ** p-value=0.0086, Welch t-test. The parapineal fails to migrate in 52% of DMSO treated *mib*^−/−^ mutant embryos (PP mean position between −15 µm to +15 µm, grey shaded area), and this proportion decreases to 19% of SU5402 treated *mib*^−/−^ mutant embryos; p-value=0.0139 on a Chi-square test and p=0.0103 on a Welch t-test on absolute value. (G) Upper panel shows a schematic of the SU5402 (or DMSO) treatment timeline (24-32 hpf) in control and *mib*^−/−^ mutant embryos. Dot plot showing the number of *Tg(dusp6:d2EGFP)* expressing parapineal cells at 32 hpf in control embryos treated with DMSO (light blue; n=20) or with 5µM SU5402 (dark blue, n=18), and in *mib*^−/−^ mutant embryos treated with DMSO (orange, n=17) or with SU5402 (yellow, n=21). The number of *Tg(dusp6:d2EGFP)+* cells is increased in DMSO treated *mib*^−/−^ mutants versus controls (p-value=0.0077) but is decreased in SU5402 treated *mib-/-*mutants compared to DMSO treated *mib-/-*mutants (p-value=0.0248 in Welch’s t-test). (H-I) Upper panel in H shows a schematic of heat shock timeline in *Tg(hsp70:ca-fgfr1)* embryos injected with *rbpj a/b* morpholinos (MO). Dot plots showing the number (H) and the mean position (I) of *gfi1ab* expressing parapineal cells in control embryos that carry (purple dots; n=55) or do not carry the *Tg(hsp70:ca-fgfr1)* transgene (light blue dots; n=54), or in *rbpja/b* MO injected embryos that carry (dark red dots; n=73) or do not carry the *Tg(hsp70:ca-fgfr1)* transgene (orange dots; n=67). In *rbpja/b* morphants expressing *Tg(hsp70:ca-fgfr1)*, the parapineal failed to migrate in 25% of embryos (n=18/73; p-value=0.0007 Welch’s t-test on absolute value and p-value=0.0232 in Chi-square test), a defect significantly higher than expected from merely adding the effects of activated receptor transgene and *rbpja/b* MO injections alone (p-value=0.0001 in Welch t-test on absolute value and p-value=0.0003 in Chi-square test). Data are representative of three (A-F, H-I) or two experiments (G)

Suboptimal SU5402 treatment had no effect on parapineal cells specification, either in *mib*^−/−^ mutant embryos or siblings (**Figure 4E**). As previously observed, we detected an increase in the number of *gfi1ab* expressing parapineal cells in DMSO treated *mib*^−/−^ embryos (18±5) compared to controls (15±2; p-value=0.0042). The number of *gfi1ab* positive cells at 48 hpf was also increased in SU5402 treated *mib*^−/−^ embryos (19±7) compared to SU5402 treated controls (15±3; p-value=0.0086); however, there was no significant difference in the number of *gfi1ab* expressing cells between DMSO treated *mib*^−/−^ embryos and SU5402 treated *mib*^−/−^ embryos (p-value=0.5687). These data support further that Notch signaling controls the number of parapineal cells independently from its function in parapineal migration and doesn’t involve Notch mediated modulation of FGF signaling.

To address the connection between Notch and FGF signaling in an alternative way, we asked whether ectopic activation of the FGF pathway could elicit a more severe phenotype in a loss-of-function context for Notch. To achieve this, we used a transgenic line that expressed a constitutively activated Fgf receptor after heat shock, *Tg(hsp70l:Xla.Fgfr1,cryaa:DsRed)* (Marques et al., 2008), in embryos injected with *rbpja/b MO* (4 to 8 ng *rbpja/b* MO); we chose this Notch context as parapineal migration defects are more modest than in *mib*^−/−^ mutants. As previously described (Roussigné et al., 2018), widespread expression of constitutively activated receptor (CA-FgfR1) prevented parapineal migration in a small number of embryos (7% of embryos, n=4/55, parapineal mean position between −15 and +15 µm; p-value=0.0011), while the parapineal consistently migrates in heat shocked control embryos not carrying the transgene (n=54/54 with n=51/54 leftwards and n=3/54 toward the right) (**Figure 4I**). In the absence of the CA-FgfR transgene, *rbpja/b* morphant embryos displayed a migration defect at low frequency (7%; n=5/67) (**Figure 4I**). However, in *rbpja/b* morphants expressing the activated receptor, the frequency of embryos in which the parapineal failed to migrate increased significantly (25% of embryos; n=18/73) (**Figures 4I,** red versus yellow and purple dots). The interaction detected between the Notch and FGF pathways in this context appears synergistic as the increase in parapineal migration defects observed in *rbpja/b* morphant embryos expressing the activated Fgf receptor is significantly higher than expected from adding the effects of activated receptor transgene and *rbpja/b* MO injections alone (p-value=0.0001 in Welch t test on absolute value and p-value=0.0003 in Chi-square test). The increased frequency of parapineal migration defects occurs in the absence of significant changes in parapineal cell-type specification (**Figures 4H**); the mean number of *gfi1ab* expressing parapineal cells did not vary significantly between heat shocked *rbpja/b* morphant embryos with or without the CA-FgfR transgene (14±6 versus 13±5; p-value=0.39).

Taken together, our loss-of-function, gain-of-function and epistasis experiments argue that the Notch pathway is required for parapineal migration, and that it acts by restricting FGF activation in parapineal cells.

## Discussion

In this study, we address how activation of the FGF pathway is restricted within parapineal cells. We show that expression of the *Tg(dusp6:d2EGFP)* reporter transgene is broader in Notch loss-of-function contexts and is reduced following global activation of the Notch pathway. These changes in FGF activation correlate with defect in parapineal migration. Epistasis experiments show that decreasing or increasing FGF signaling can respectively rescue or aggravate parapineal migration defects in Notch loss-of-function contexts. We conclude that the Notch pathway participates in restricting the activation of FGF signaling to few cells of the parapineal cluster, thus promoting its migration.

### Notch effects on parapineal cell specification can be uncoupled from migration

Studies addressing the link between cell specification and migration are rare and, as such, our knowledge of whether and how these two processes are coordinated is limited. In the lateral line primordium (LLP) model, proper morphogenesis of the future neuromast at the trailing edge of the primordium is required for migration (Durdu et al., 2014; Kozlovskaja-Gumbrienė et al., 2017; Lecaudey et al., 2008; Nechiporuk and Raible, 2008). In embryos treated with LY411575 from 22 to 32 hpf, the number of *gfi1ab* and *sox1a* expressing parapineal cells is strongly increased but neither parapineal migration nor *Tg(dusp6:d2EGFP)*+ expression are affected. Therefore, our data reveals that cell-type specification and migration can be uncoupled in the parapineal. Our results resemble previous observations describing uncoupling of specification and migration of cardiac cells during in heart development (Davidson et al., 2005).

Notch mediated cell-cell communication is well described for its role on cell fate and progenitors maintenance (Cau and Blader, 2009). In neural tissues or in the pancreas, for instance, inhibition of the Notch signaling pathway causes premature differentiation of the progenitor cells into mature differentiated cells (Li et al., 2015). In loss or gain of function for Notch, the number of *gfi1ab* and *sox1a* expressing parapineal cells is affected. However, in both contexts the parapineal rosette is formed and a normal number of *tbx2b* expressing parapineal cells is detected. This suggests that Notch signaling acts downstream of Tbx2b, and probably controls the transition from *tbx2b* expressing putative progenitors to differentiated parapineal cells expressing *sox1a/gfi1ab*.

### Notch acts upstream of FGF signaling to promote parapineal migration

In most models describing cross-talk between the Notch and FGF pathways, Notch signaling is described to act downstream of the FGF pathway. In the zebrafish LLP, for instance, ectopic activation of Notch signaling can rescue the formation of neuromast rosettes in absence of FGF pathway activity, indicating that Notch signaling is required downstream of Fgf signals to promote apical constriction and rosette morphogenesis (Kozlovskaja-Gumbrienė et al., 2017). Notch signaling is also described to be a downstream effector of the FGF pathway for epithelial proliferation in the pancreas in mammalian embryos (Hart et al., 2003). Our epistasis experiments suggest that the Notch pathway acts upstream of FGF signaling in parapineal cells to restrict FGF pathway activation and promote migration. Therefore, the crosstalk between the FGF and Notch pathway appears highly context specific.

### How can Notch restrict the activation of FGF pathway?

Notch signaling has previously been implicated in the migration of epithelial cells sheets (Riahi et al., 2015), trachea cells in *Drosophila* (Ghabrial and Krasnow, 2006; Ikeya and Hayashi, 1999) and in developing vertebrate blood vessels (Siekmann and Lawson, 2007b). Interestingly, in the latter two examples, Notch-Delta signaling contributes to the selection of the tip cells by restricting the ability of followers cells to activate RTK signaling although the molecular mechanisms are not clear yet (Ghabrial and Krasnow, 2006; Ikeya and Hayashi, 1999; Siekmann and Lawson, 2007b). In all these models, migrating cells remains attached to the bulk of the tissue. The parapineal is thus the first described model of an isolated cluster of migrating cells in which Notch signaling modulates RTK signaling to define leading cells.

Our data indicate that the Notch pathway promotes migration by restricting the activation of FGF signaling to a few parapineal cells, but how Notch signaling could modulate FGF pathway activity remains an open question. The fact that some *rbpja/b* morphants display parapineal migration defects similar to *mib*^−/−^ mutants strongly supports a role for the canonical Notch pathway rather than a Notch independent role of the Mindbomb ubiquitin ligase in parapineal migration, although we cannot exclude that Mindbomb also regulates migration independently of the Notch pathway, for instance through modulation of Rac1 (Mizoguchi et al., 2017). As all PP cells are competent to migrate and to activate the FGF pathway (Concha et al., 2003; Roussigné et al., 2018), a parsimonious mechanism would be that lateral inhibition based cell-cell communication between parapineal cells within the rosette would modulate the capacity of a cell to activate/maintain or to inhibit FGF pathway activity. As the ectopic expression of the constitutive activated FGF receptor (CA-FgfR1) does not completely block parapineal migration in a wild-type background and can rescue migration in *fgf8*^−/−^ mutants (Roussigné et al., 2018), it appears that the restriction mechanism acts downstream of the receptor rather than at the level of receptor gene expression. As described for the *C. elegans* vulva (Berset et al., 2001; Yoo et al., 2004), Notch signaling could directly promote the transcription of Ras/MAPK pathway inhibitors.

### Time window of Notch requirement

The fact that ectopic Notch signaling at late stages (26-28 hpf) can efficiently block migration argues for a role of Notch at the time when the parapineal initiates its migration. If Notch signaling is indeed required just before migration, it is unclear why LY411575 treatment (22 to 32 hpf) does not affect migration while it is able to trigger an increase in the number of *gfi1ab* and *sox1a* positive parapineal cells.

Classically, Notch signaling is thought to act through a lateral inhibition mechanism leading to a Notch-ON or Notch-OFF outcome between neighbouring cells. Recent work, however, suggests that Notch acts in a level-dependent manner rather than in an all-or-nothing mode (Ninov et al., 2012). In light of this, the differential effect of LY411575 on specification and migration of parapineal cells could reflect a different requirement in Notch signaling threshold, with a lower threshold of Notch signaling being required for migration and a higher one being required to control fate specification of parapineal cells. If LY411575 does not completely block Notch signaling, then residual Notch activity could be sufficient to promote the restriction of FGF activity and parapineal migration while not to limit the specification of *gfi1ab* and *sox1a* positive cells. Consistent with the hypothesis that LY411575 treatment might be partially effective, we observed that the average number of *Tg(dusp6:d2EGFP)*+ cells increases slightly in LY411575 treated embryos from 22 to 32 hpf, without reaching statistical significance. Given the high variability observed in the number of *Tg(dusp6:d2EGFP)*+ cells in wild-type contexts, we might expect that parapineal migration is robust enough to tolerate a moderate increase in the number of *Tg(dusp6:d2EGFP)+* cells.

### Synergistic or parallel role of the Nodal and Notch pathways in restricting FGF pathway activation

The Notch signaling pathway has previously been implicated in regulating the expression of Nodal/TGF-β signal around the node and subsequently in the left lateral plate mesoderm (LPM) (Krebs et al., 2003; Raya et al., 2003). In zebrafish, expression of a *nodal* related gene (*ndr3/southpaw*) in the left LPM is required for the later expression of a second *nodal* gene (*ndr2*/*cyclops*) in the left epithalamus, which is required for left-biasing parapineal migration (Concha et al., 2000; Liang et al., 2000; Regan et al., 2009). In our Notch loss-of-function contexts, the Nodal pathway is activated on both sides of the epithalamus. This requirement for Notch signaling in unilateral activation of the Nodal pathway in the epithalamus is consistent with an early role of Notch signaling in establishing the initial left right asymmetry andexplains the partial randomization of parapineal migration we observe in contexts of early Notch loss-of-function.

Our previous results indicate that Nodal signaling contributes to restricting FGF pathway activation, as well as biasing it to the left (Roussigné et al., 2018). Here, we show that the restriction of FGF activity requires a functional Notch pathway. As mentioned above, the Nodal pathway is bilateral in the epithalamus of embryos with compromised Notch signaling. However, it is unlikely that the role of Notch in restricting FGF pathway activation depend on its ability to control Nodal signaling. Indeed, in other contexts of bilateral Nodal signaling, such as in embryos injected with morpholinos against *notail*, we have shown that FGF pathway activation ultimately becomes restricted to a few parapineal cells (Roussigné et al., 2018); *Tg(dusp6:d2EGFP)* reporter expression in this context is no longer left lateralized, which correlates with the parapineal migrating either to the left or the right. In absence of Nodal signaling, as in *spaw* morphants, *Tg(dusp6:d2EGFP)* expression is generally less restricted within the parapineal and this correlates with delayed parapineal migration, indicating that the restriction of FGF activity is influenced by Nodal signaling as well as by Notch pathway. How these two pathways interact to restrict the activation of FGF signaling is not known and future investigations well be needed to address whether Nodal and Notch pathway act in a synergic way or in parallel to restrict the FGF pathway.

## Conclusion

Our study shows that Notch signalling is required for parapineal migration through its capacity to trigger cell state differences in FGF signaling and to restrict FGF activity to a few leading cells. As the function and cross-regulation of FGF and Notch pathways might be conserved during the migration of invading cancerous cells, our results could help to understand these pathway interaction during metastasis.

## Materials and Methods

### Fish lines

Embryos were raised and staged according to standard protocols (Westerfield, 2000). Embryos homozygous for *midbomb* (*mib*^*ta52b*^) (Itoh et al., 2003) and *fgf8* mutations (*fgf8* ^*ti282a*^ */ acerebellar / ace*; (Reifers et al., 1998)) were obtained by inter-crossing heterozygous carriers. Carriers of the *fgf8*^*ti282a*^ allele were identified by PCR as described previously (Roussigné et al., 2018). *mib*^*ta52b+/−*^ carriers were identified by PCR genotyping using primers 5’-GGTGTGTCTGGATCGTCTGAAGAAC-3’ and 5’-GATGGATGTGGTAACACTGATGACTC-3’ followed by enzymatic digestion with NlaIII. *Tg(hsp70:Gal4)*^*kca4*^ and *Tg(UAS:myc-Notch1a-intra)*^*kca3*^ transgenic lines have been described previously (Scheer et al., 2001; Scheer and Campos-Ortega, 1999) and identified by PCR genotyping following ZIRC genotyping protocols. Embryos carrying the *Tg(hsp70:ca-fgfr1; cryaa:DsRed)*^*pd3*^ transgene (Marques et al., 2008; Neilson and Friesel, 1996) were identified by the presence of DsRed expression in the lens from 48 hpf or, by PCR at earlier stages as described previously (Gonzalez-Quevedo et al., 2010). *Tg(dusp6:d2EGFP)*^*pt6*^ (Molina et al., 2007) lines were used as reporters for FGF pathway activity. Embryos were fixed overnight at 4°C in 4% paraformaldehyde/1xPBS, after which they were dehydrated through an ethanol series and stored at −20°C until use.

### Ethics statement

Fish were handled in a facility certified by the French Ministry of Agriculture (approval number A3155510). The project has received an agreement number APAFIS#3653-2016011512005922. Anaesthesia and euthanasia procedures were performed in Tricaine Methanesulfonate (MS222) solutions as recommended for zebrafish (0,16mg/ml for anaesthesia, 0,30 mg/ml for euthanasia). All efforts were made to minimize the number of animals used and their suffering, in accordance with the guidelines from the European directive on the protection of animals used for scientific purposes (2010/63/UE) and the guiding principles from the French Decret 2013-118.

### LY411575 treatment

Embryos collected from *Tg(dusp6:d2GFP)*^*pt6*^ carriers outcrossed with wild-type fish or with heterozygous *mib*^*ta52b*^ were dechorionated and treated from 8 to 22 hpf or 22 to 32 hpf with 30 µM, 100 µM or 200µM of LY411575 (MedChem; (Rothenaigner et al., 2011)); control embryos were treated with an equal volume of DMSO diluted in E3 medium. *Tg(dusp6:d2GFP)* expressing embryos were incubated in LY411575 at 28°C and fixed at 32 hpf for immune-staining against EGFP; sibling embryos, not carrying the *Tg(dusp6:d2GFP)* transgene, were fixed at indicated time (32 hpf, 36 hpf or 48 hpf) for in situ against different parapineal markers.

### SU5402 drug treatment

Embryos collected from in-crosses between *mib*^*ta52b*^ and *Tg(dusp6:d2EGFP); mib*^*ta52b*^ mutants were dechorionated and treated with 5µM SU5402 (Calbiochem; (Mohammadi et al., 1997)) by diluting a 10 mM DMSO based stock solution in E3 medium; control embryos were treated with an equal volume of DMSO diluted in E3 medium. All embryos were incubated in SU5402 at 28°C from 24 hpf to 32 hpf. *Tg(dusp6:d2EGFP)* embryos were fixed at 32 hpf to analyze the d2GFP expression pattern in both *mib*^*ta52b*^ mutants and sibling embryos; the remaining half of embryos were fixed at 48 hpf to analyze parapineal migration.

### Morpholino injections

Morpholino oligonucleotides targeting both *rbpja /su(H)1* and *rbpjb/su(H)2* (Echeverri and Oates, 2007) were dissolved in water at 3 mM. The resulting stock solution was diluted to working concentrations (0,3 mM, 2.5ng/nl) in water and Phenol Red before injection of 1,5 nl (4ng) or 4nl (10ng) into embryos at the 1 cell stage. Embryos were subsequently fixed and processed for ISH or antibody labelling.

### Heat shock procedure

Ectopic expression of the intracellular domain of Notch receptor (NICD) was induced in *Tg(hs70:gal4); Tg(uas:notch1a-intra-myc)* double transgenic embryos by incubating them at 39°C for 45 minutes starting at different time points (22 hpf, 24 hpf, 26 hpf, 28 hpf or 32 hpf). Embryos were then incubated at 28.5 °C and fixed at 36 hpf to analyze *Tg(dusp6:d2EGFP)* expression or at 48 hpf for *in situ* against indicated parapineal markers. Ectopic expression of CA-FgfR1 was induced in *Tg(hsp70:ca-FgfR1; cryaa:DsRed)*^*pd3*^ heterozygote embryos by performing a first heat shock at 25-26 hpf (39°C, 45 minutes) and a second short heat shock (39°C, 15 min) 3 hours later (28-29 hpf) in order to cover the entire period of parapineal migration.

### In situ hybridization and immunostaining

Embryos were fixed overnight at 4°C in BT-FIX (Westerfield, 2000) after which they were dehydrated through ethanol series and stored at −20°C until use. In situ hybridizations were performed as described previously (Roussigné et al., 2018), using antisense DIG labelled probes for *gfi1ab* (Dufourcq et al., 2004), *sox1a* (Clanton et al., 2013), *pitx2c* (Essner et al., 2000) and *tbx2b* (Snelson et al., 2008). In situ hybridizations were completed using Fast Red (from Roche or Sigma Aldrich) as an alkaline phosphatase substrate. Immuno-stainings were performed in PBS containing 0,5% triton using anti-GFP (1/1000, Torrey Pines Biolabs) and Alexa 488 or Alexa 555-conjugated goat anti-rabbit IgG (1/1000, Molecular Probes). For nuclear staining, embryos were incubated in ToPro-3 (1/1000, Molecular Probes) for 1h as previously described (Roussigné et al., 2018).

### Image acquisition

Confocal images of fixed embryos were acquired on an upright Leica SP8 confocal microscope using the resonant fast mode and either an oil x63 (aperture 1.4) or x20 (aperture 1.4) objective. Confocal stacks were analyzed using ImageJ software. Figures were prepared using Adobe Photoshop software.

### Quantification of the number and position of *Tg(dusp6:d2EGFP)* positive parapineal cells

The position and number of parapineal cells positive for the *Tg(dusp6:d2GFP)* transgene were analyzed using ROI Manager tool on ImageJ software as previously described (Roussigné et al., 2018). The mean intensity of the d2EGFP staining was quantified in an area corresponding to the cell nucleus and the same intensity threshold was used in the different experimental contexts to determine if a cell was *Tg(dusp6:d2EGFP)* positive or not. The total number of parapineal cells was estimated by counting nuclei in the parapineal rosettes using Topro-3 nuclear staining. For each parapineal cell, we calculate its x and y position relative to the center of the parapineal (calculated as the mean of x and y positions of all parapineal cells).

### Quantification of the number and position of *gfi1ab, sox1a, tbx2b* positive parapineal cells

The position and number of *gfi1ab, sox1a* or *tbx2b* positive parapineal cells were analyzed using the Multipoint tool on ImageJ software and determined as the centre of the cell nucleus detected with the Topro-3 nuclear staining. The position of each parapineal cell was measured relative to the brain midline (reference origin =0) as determined by a line passing along the lumen of the epiphysis. For each embryo, we calculated the number of labelled parapineal cells and their mean position. To avoid any bias, data in Figure 4E-F were analysed blind for DMSO versus SU5402 treatment and data describing LY411575 treatment (Figure 2I-N, Figure 2–figure supplement 1) were analysed blind for *mib* genotypes (*mib*+/− heterozygotes versus +/+ controls).

### Statistical analysis

Statistical comparisons of datasets were performed using R Studio or GraphPad Prism software. For each dataset, we tested the assumption of normality with Shapiro-Wilks tests and variance homogeneity with F tests. As datasets on the number of parapineal cells were usually normal and often of unequal variances, they were compared using unpaired Welch t-tests; means (± SEM) are indicated as horizontal bars on dot plots. Data on parapineal mean position usually did not distribute normally (because of few embryos with the parapineal on the right) and were compared using the Wilcoxon rank sum non-parametric tests; we also compared parapineal mean position datasets with Welch t-tests using absolute values to discriminate between a defect in laterality (i.e. left-right randomization of parapineal migration) and a defect in migration *per se* (i.e. distance from the midline). Most data are representative of at least two and more often three independent experiments and the number of biological replicates is mentioned in the figure legends for each graph. Means (± SD) are indicated in the text.

## Acknowledgements

We are grateful to all members of the Blader lab, especially Elise Cau and Julie Batut for critical reading of the manuscript and Aurélie Quillien for insightful discussions. We also thank Eric Theveneau and Matthias Carl for critical reading of the manuscript, as well as Xiaobo Wang and Steve W. Wilson for their input. We thank Brice Ronsin from the Toulouse RIO Imaging platform and Aurore Laire for fish care. This work was supported by the Centre National de la Recherche Scientifique (CNRS), the Institut National de la Santé et de la Recherche Médicale (INSERM), Université de Toulouse III (UPS), the Fondation pour la Recherche Médicale (FRM; DEQ20131029166), the Fondation ARC (PJA 20131200173), the Agence nationale de la recherche (ANR-16-CE13-0013-01) and the China Scholarship Council (CSC) for PhD funding for Lu Wei.

---

**Figure 1–figure supplement 1.**
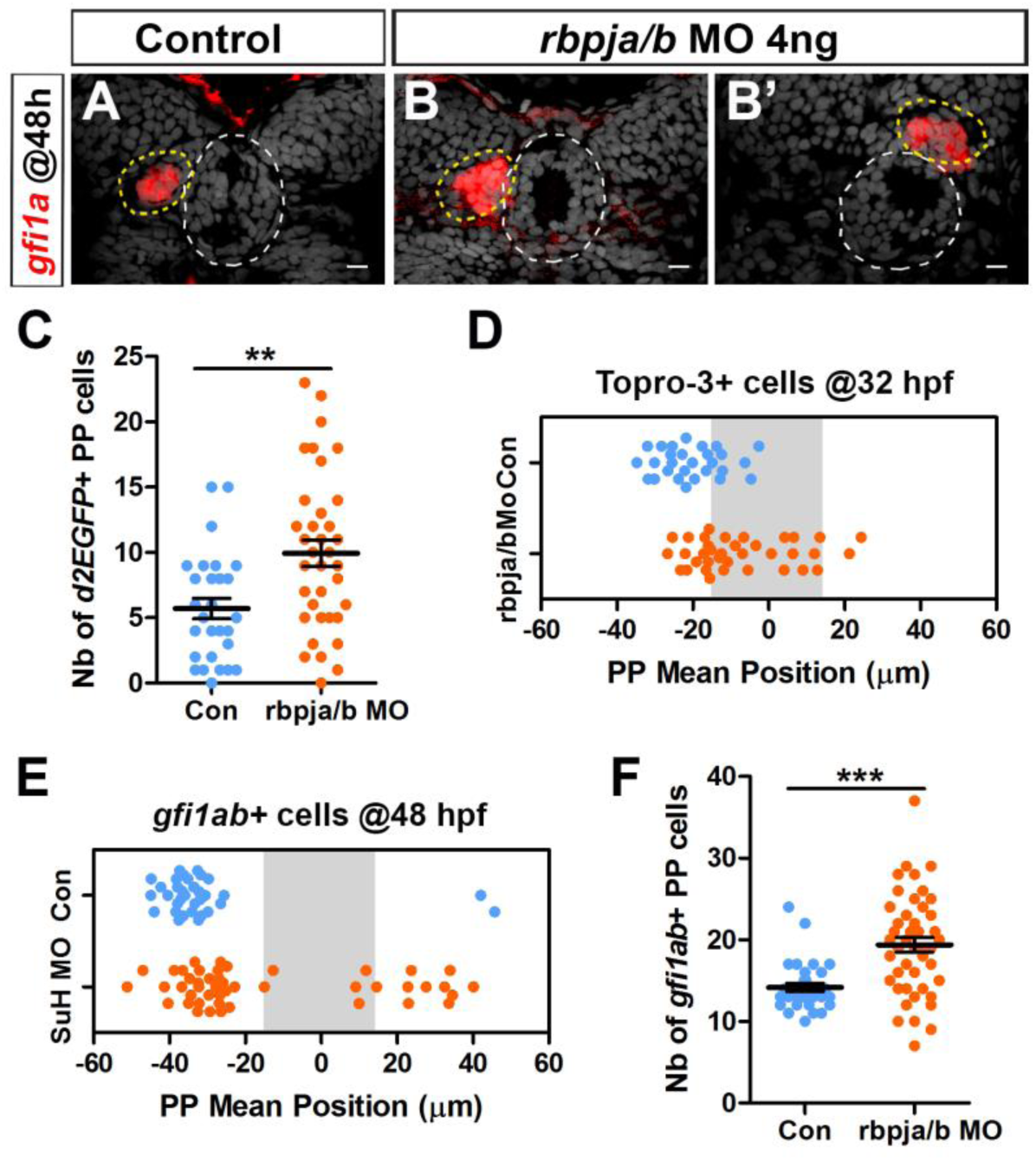
rbpja/b morphants phenocopy mindbomb mutants. (A-B’) Confocal sections showing the expression of *gfi1ab* (red) at 48 hpf in control embryos (A, n=33) and in 2 representative embryos injected with 4 ng *of rbpja/b* MO (B, B’); sections are merged with nuclear staining (grey) to visualize the epiphysis (white circle) and parapineal (yellow circle). Embryo view is dorsal, anterior is up; scale bar=10 µm. (C-D) Dot plots showing the number of *Tg(dusp6:d2EGFP)* positive parapineal cells (C) and the mean position of parapineal cells visualized by nuclei staining (Topro-3) (D) at 32 hpf in embryos injected with 4 ng *rbpja/b* MO (orange dots; n=36) or in controls (blue dots; n=28). (E-F) Dot plots showing the mean position (E) or the number (F) of *gfi1ab* expressing parapineal cells in non-injected controls (blue dots, n=33) or in *rbpja/b* MO injected embryos (orange dots, n=46) at 48 hpf. Both the migration (distance from the midline; p-value=0.0001 in Welch t-test on absolute value) and laterality (left orientation of migration; p-value=0.0002 on Wilcoxon test) are affected in *rbpja/b* morphants (E). The number of *gfi1ab+* cells is also increased in *rbpja/b* morphants (F, *** p-value<0.0001; Welch t-test). Data are representative of three (A-B’, E-F) or two experiments (C-D).

**Figure 2–figure supplement 1.**
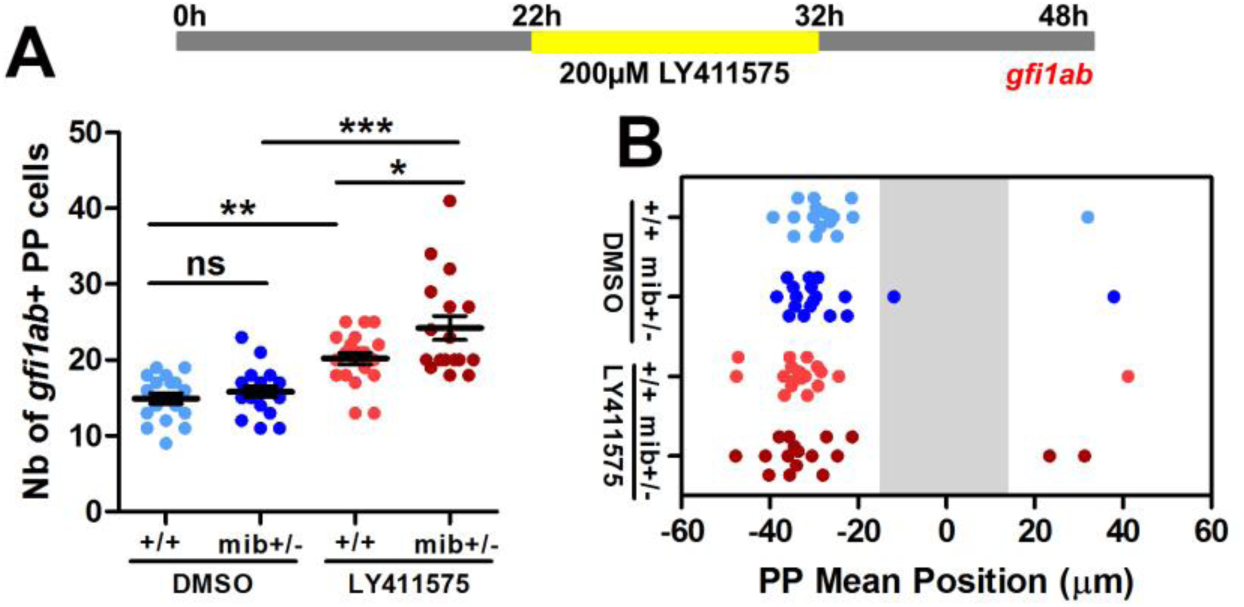
Effect of treatment with high dose of γ-secretase inhibitor from 22 to 32 hpf on the specification and migration of parapineal cells. (A, B) Upper panel shows a schematic of LY411575 treatment timeline: wild-type (+/+) or *mib*^+/−^ heterozygote embryos were treated from 22 to 32 hpf with 200 µM of LY411575. Dot plots show the number (A) and the mean position (B) of *gfi1ab* expressing cells at 48 hpf in the parapineal of DMSO treated controls (dark blue dots; n=18), DMSO treated *mib*^+/−^ heterozygotes (light blue dots; n=18), LY411575 treated (dark red dots; n=20) and LY411575 treated *mib*^+/−^ heterozygotes (light red dots, n=17); dots represent individual embryos. *** p-value<0.0001, ** p-value<0.01, * p-value<0.05, Welch t-test and Wilcoxon test; Data corresponds to one experiment.

**Figure 2–figure supplement 2.**
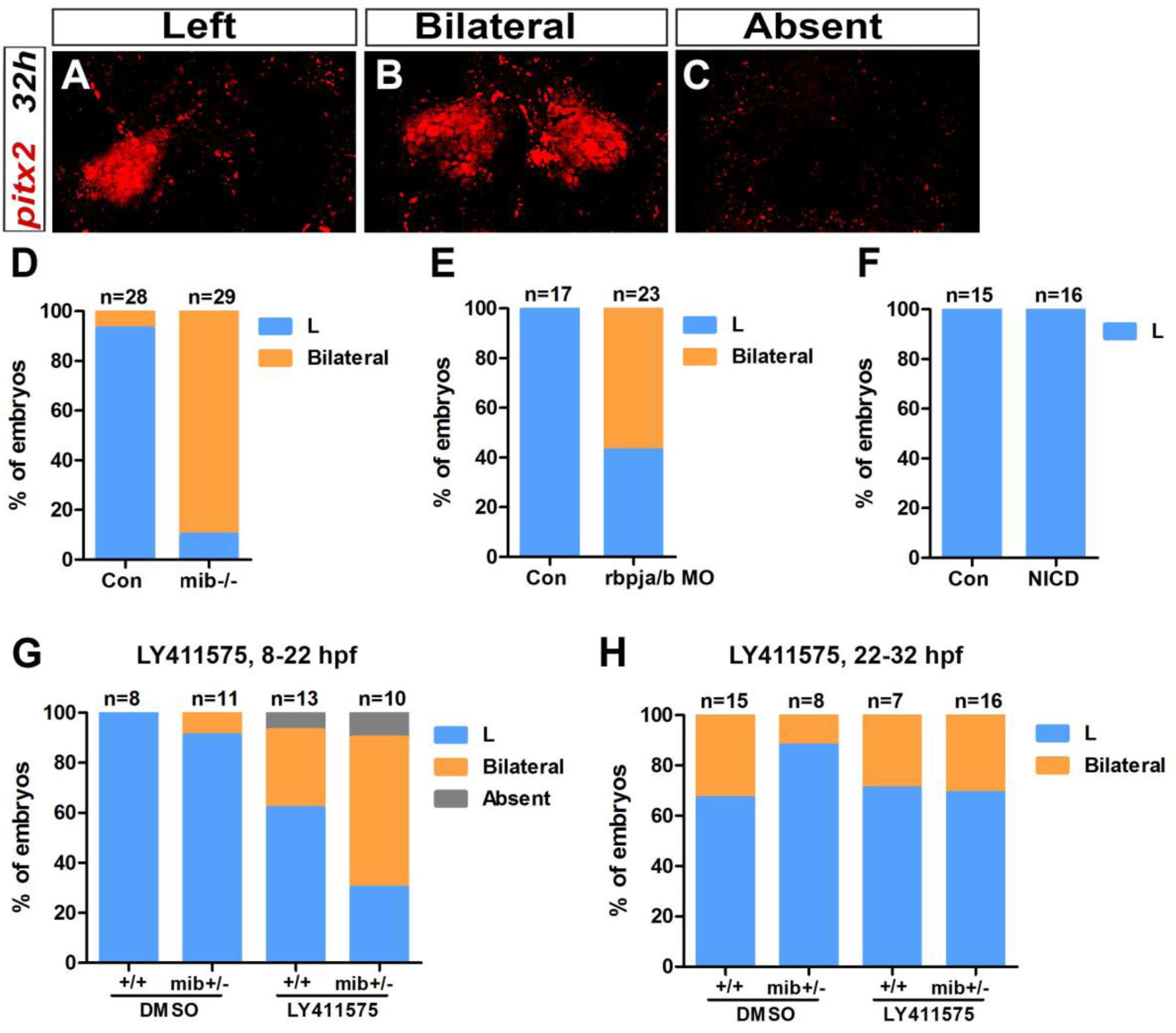
Early Notch loss-of-function results in bilateral Nodal pathway activation in the epithalamus. (A-C) Confocal sections showing Left (A), Bilateral (B) or Absent (C) *pitx2c* expression in the epithalamus at 32 hpf (red). Embryo view is dorsal, anterior is up; scale bar=10 µm. (D-H) Histogram showing the percentage of embryos with Left (L, blue), Bilateral (orange) and Absent (grey) *pitx2c* expression in *mib*^−/−^ mutant embryos (D), in *rbpja/b* morphant (MO) embryos (E), in embryos expressing NICD (Notch Intra-cellular Domain) (F) and in embryos treated with 30µM of LY411575 from 22 to 32 hpf (G) or from 8 to 22 hpf (H). Genetic background, treatment and embryos numbers are indicated below each bar; Con: control sibling embryos. Data are representative of two experiments (D-F) or one experiment (E, G, H).

**Figure S3–figure supplement 1.**
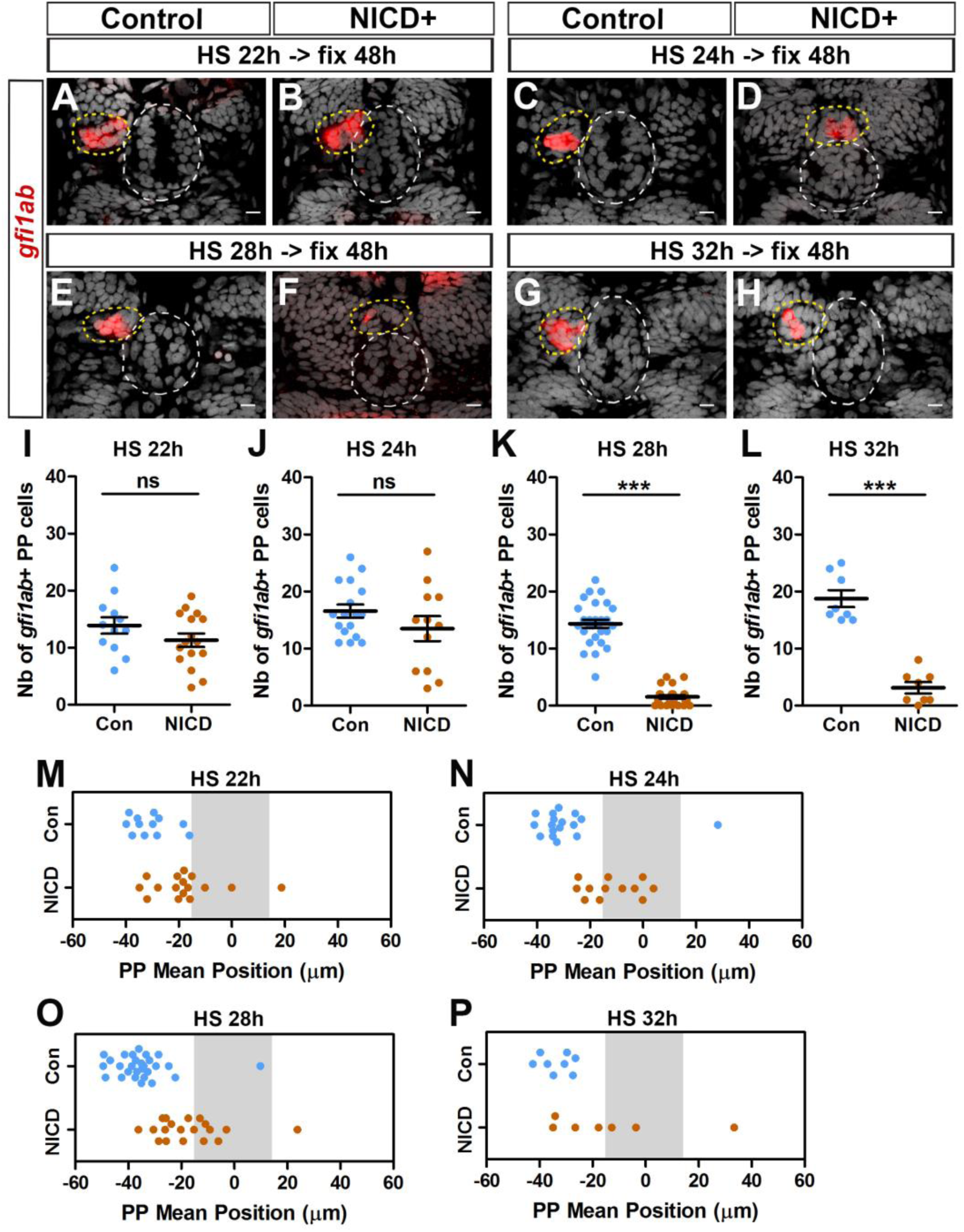
Effect of activation of the Notch pathway at 22, 24, 28 and 32 hpf on the migration and specification of parapineal cells. (A-H) Confocal maximum projections showing the expression of *gfi1ab* (red) at 48 hpf in control embryos (A, n=27; C, n=17; E, n=28; G, n=8) or in *Tg(hsp70l:Gal4);Tg(UAS:myc-Notch1a-intra)* double heterozygous embryos (B, n=16; D, n=12; F, n=26; H, n=8) following heat-shock (HS) induction at 22 hpf (A, B), 24 hpf (C, D), 28 hpf (E, F), 32 hpf (G, H); images are merged with nuclear staining (grey) to visualize the epiphysis (white circle) and parapineal (yellow circle). Embryo view is dorsal, anterior is up; scale bar=10 µm. (I-P) Dot Plots showing the quantification of the number (I, J, K, L) and the mean position (M-P) of *gfi1ab* expressing parapineal (PP) cells in control (blue dots) or in *NICD+* embryos (orange dots) at 48 hpf, following heat-shock (HS) induction at 22 hpf (I, M), 24 hpf (J, N), 28 hpf (K, O), 32 hpf (L, P). The number of *gfi1ab+* parapineal cells was significantly decreased in embryos expressing NICD from 28 and 32 hpf but not from 22 and 24 hpf; mean ± SEM is indicated on dot plots I-L; *** p-value<0.0001 in Welch t-test and Wilcoxon test. Parapineal migration was affected in embryos expressing NICD from 22, 24 or 28 hpf; p-value=0.0018 (M), <0.0001 (N), <0.0001 (O) in Welch t-test in absolute values; defects were variable but also significant in embryos expressing NICD from 32 hpf, p=0.0369 (P). Data are representative of three (HS@28 hpf; E-F, K, O) or two experiments (HS@24 hpf; C-D, J, N); data shown in A-B, I, M and G-H, L, P (HS@22 hpf and 32 hpf) correspond to one experiment.

